# Vagal nerve stimulation triggers widespread responses and alters large-scale functional connectivity in the rat brain

**DOI:** 10.1101/200220

**Authors:** Jiayue Cao, Kun-Han Lu, Terry L. Powley, Zhongming Liu

## Abstract

Vagus nerve stimulation (VNS) is a therapy for epilepsy and depression. However, its efficacy varies and its mechanism remains unclear. Prior studies have used functional magnetic resonance imaging (fMRI) to map brain activations with VNS in human brains, but have reported inconsistent findings. The source of inconsistency is likely attributable to the complex temporal characteristics of VNS-evoked fMRI responses that cannot be fully explained by simplified response models in the conventional model-based analysis for activation mapping. To address this issue, we acquired 7-Tesla blood oxygenation level dependent fMRI data from anesthetized Sprague–Dawley rats receiving electrical stimulation at the left cervical vagus nerve. Using spatially independent component analysis, we identified 20 functional brain networks and detected the network-wise activations with VNS in a data-driven manner. Our results showed that VNS activated 15 out of 20 brain networks, and the activated regions covered >76% of the brain volume. The time course of the evoked response was complex and distinct across regions and networks. In addition, VNS altered the strengths and patterns of correlations among brain networks relative to those in the resting state. The most notable changes in network-network interactions were related to the limbic system. Together, such profound and widespread effects of VNS may underlie its unique potential for a wide range of therapeutics to relieve central or peripheral conditions.

## Introduction

Since the 1800s (1, 2), vagus nerve stimulation (VNS) has been studied as a potential way to treat various diseases, including epilepsy, depression, tinnitus, Alzheimer’s Disease, and obesity (3-8). The therapeutic benefits apparently depend on the effects of VNS on the central neural system (CNS) mediated through neuroelectrical or neurochemical signaling (9). Studies have been conducted to evaluate the CNS responses to VNS with neural imaging or recording techniques. For example, invasive recordings of unit activity or field potentials have shown VNS-evoked neuronal responses in the nucleus of solitary tract (10), the locus coeruleus (11), and the hippocampus (12). These techniques offer high neuronal specificity but only cover spatially confined targets. In contrast, electroencephalogram (EEG) has been used to reveal VNS-induced synchronization or desynchronization of neural oscillations in the macroscopic scale (13, 14), while being severely limited by its spatial resolution and specificity as well as its inability to detect activities from deep brain structures. However, sub-cortical regions are of interest for VNS, because the vagus nerves convey signals to the brain through polysynaptic neural pathways by first projecting to the brainstem, then subcortical areas, and lastly the cortex (9, 15).

Complementary to conventional electrophysiological approaches, functional neuroimaging allows characterizing the effects of VNS throughout the brain volume. Using positron emission tomography (PET) or single-photon emission computerized tomography (SPECT), prior studies have reported VNS-evoked responses in the thalamus, hippocampus, amygdala, inferior cerebellum, and cingulate cortex (16-19); but these techniques are unable to capture the dynamics of the responses due to their poor temporal resolution. In this regard, functional magnetic resonance imaging (fMRI) is more favorable because it offers balanced and higher spatial and temporal resolution. Previous human VNS-fMRI studies have reported VNS-evoked blood oxygenation level dependent (BOLD) responses in the thalamus, hypothalamus, prefrontal cortex, amygdala, and hippocampal formation (20-24). However, the reported activation patterns are not always consistent (25), sometimes highlighting activations in different regions or even opposite responses in the same regions (20, 22). What underlies this inconsistency might explain the varying efficacy of VNS in treatment of individual patients, or might be attributed to the analysis methods for activation mapping (25). Therefore, it is desirable to explore and evaluate various methodological choices in the fMRI data analysis, in order to properly interpret the VNS evoked activations for understanding the implications of VNS to neurological disorders.

Functional MRI not only localizes the CNS responses of VNS (25), it also reveals the patterns and dynamics of functional networks during VNS, which helps to characterize the network basis of VNS-based therapetutics. Findings from prior studies have shown that the therapeutic or behavioral effects of VNS may be compromised, when the underlying neuronal circuit is disrupted in terms of its critical node or receptor. For example, given a lesion in the locus coeruleus, VNS fails to suppress epilepsy (26); given a blockade of the muscarinic receptor, VNS fails to promote perceptual learning (27). However, how VNS affects the patterns of interactions among regions or networks (or functional connectivity) has rarely been addressed (28), even though fMRI has become the primary tool for studying functional connectivity (29, 30).

In this study, we aimed to address the BOLD effects of VNS in the rat brain. The use of a rat model mitigated the inter-subject variation in genetics, gender, age, weight, and health conditions. It provided a well-controlled setting for us to compare different analysis methods for mapping the activations with VNS. Specifically, we used the independent component analysis (ICA) to identify brain networks, and then used a data-driven analysis to detect the VNS-evoked activation separately for each network, as opposed to each voxel or region. In addition to the activation mapping, we also evaluated the effects of VNS on network-network interactions, against the baseline of intrinsic interactions in the resting state. As such, we attempted to address the effects of VNS on the brain from the perspectives of both regional activity and inter-regional functional connectivity.

## Methods and Materials

### Subjects

A total of 17 Sprague–Dawley rats (male, weight: 250-350g; Envigo RMS, Indianapolis, IN) were studied according to a protocol approved by the Purdue Animal Care and Use Committee (PACUC) and the Laboratory Animal Program (LAP). Of the 17 animals, seven animals were used for VNS-fMRI experiments; ten animals were used for resting state fMRI. All animals were housed in a strictly controlled environment (temperature 21±1°C and 12 h light-dark cycle, lights on at 6:00 AM, lights off at 6:00 PM).

### Animal preparation

For the VNS-fMRI experiments, each animal was initially anesthetized with 5% isoflurane and maintained with continuous administration of 2-3% isoflurane mixed with oxygen and a bolus of analgesic (Rimadyl, 5 mg/Kg, Zoetis) administrated subcutaneously. After a toe-pinch check for adequate anesthesia, a 2-3cm midline incision was made starting at the jawline and moving caudally. The left cervical vagus nerve was exposed and isolated after removing the surrounding tissue. A bipolar cuff electrode (Microprobes, made of platinum and with 1mm between contacts) was wrapped around the exposed vagus nerve. For resting-state fMRI experiments, animals were anesthetized with the same dose of anesthesia without the surgical procedures.

After the acute electrode implantation (for VNS-fMRI) or the initial anesthetization (for resting-state fMRI), each animal was moved to the small-animal horizontal MRI system (BioSpec 70/30, Bruker). The animal’s head was constrained with a customized head restrainer. A bolus of dexdomitor (Zoetis, 7.5 µg/Kg for animals gone through electrode implantation, 15 µg/Kg for animals without surgery) was administrated subcutaneously. About 15-20 mins after the bolus injection, dexdomitor was continuously and subcutaneously infused at 15 µg/Kg/h; the dose was increased every hour as needed (31). In the meanwhile, isoflurane was administered through a nose cone, with a reduced concention of 0.1-0.5% mixed with oxygen. Throughout the experiment, both the dexdomitor infusion rate and the isoflurane dose were adjusted to maintain a stable physiological condition with the respiration rate between 40 and 70 times per min and the heart rate between 250 and 350 beats per min. The heart and respiration rates were monitored by using a small-animal physiological monitoring system (Kent Scientific). The animal’s body temperature was maintained at 37 ± 0.5 °C using an animal-heating system. The oxygen saturation level (SpO_2_) was maintained above 96%.

### Vagus nerve stimulation

The bipolar cuff electrode was connected to a current stimulator (model 2200, A-M system) placed outside of the MRI room through a twisted-pair of copper wire. Stimulation current was delivered in 10s-ON-50s-OFF cycles. When it was ON, biphasic square pulses (width: 0.1 ms; amplitude: 1.0 mA; frequency: 10 Hz) were delivered. Each fMRI session included ten ON/OFF cycles. A resting (stimulation-free) period of at least one minute was given between sessions. Up to 4 sessions were scanned for each animal. Fig 1 illustrates the VNS paradigm.

**Fig 1.**
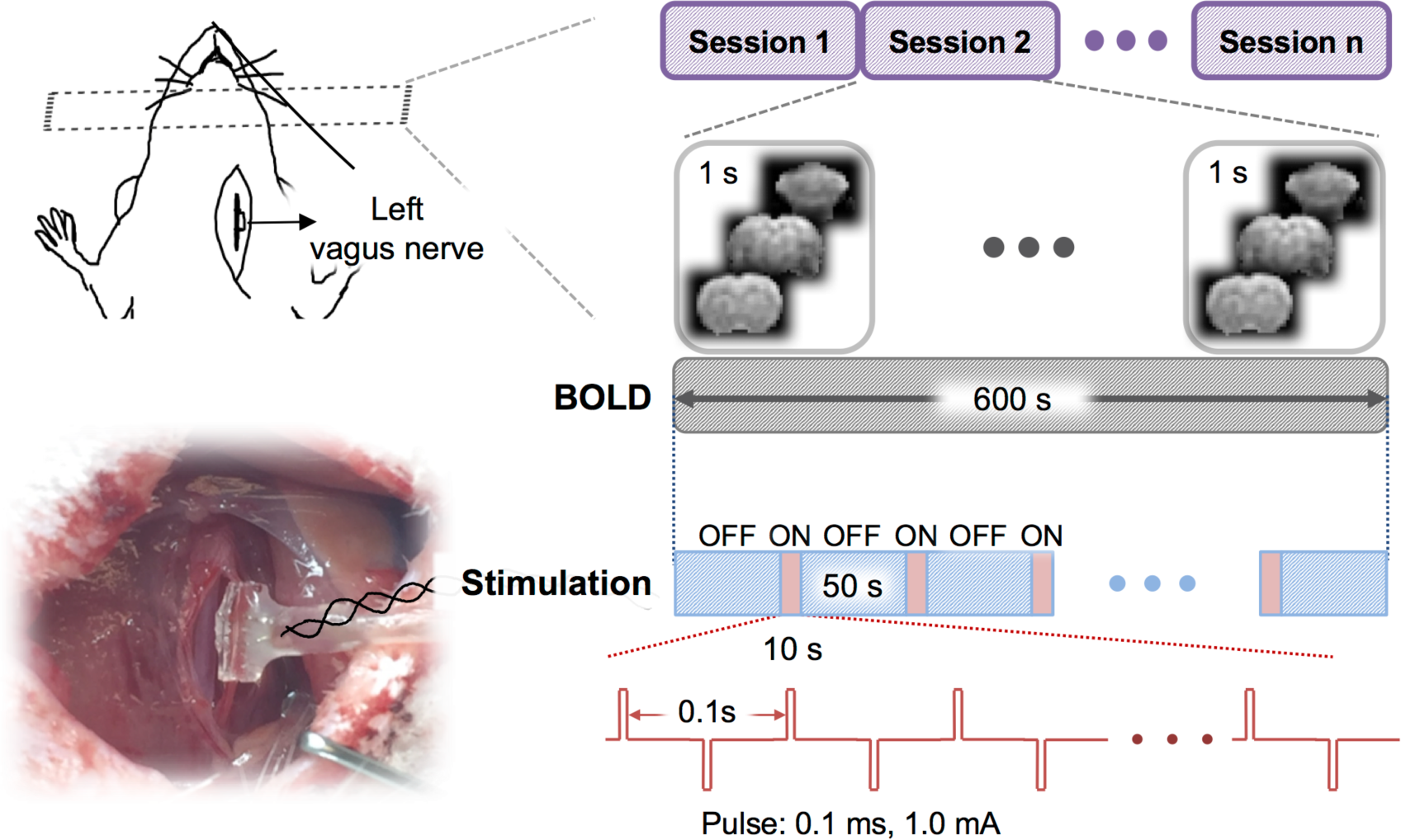
Experimental design for fMRI during VNS. Each rat was stimulated at the left cervical vagus through a cuff electrode implanted in an acute surgery. Biphasic current pulses were delivered during a 10s “ON” period alternating with a 50s “OFF” period for 10 cycles. With this block design, the rat was scanned for fMRI with a repetition time of 1s.

### MRI and fMRI

MRI data were acquired with a 7-T small-animal MRI system (BioSpec 70/30, Bruker) equipped with a volume transmitter coil (86 mm inner diameter) and a 4-channel surface receiver array. After the localizer scans, T_2_-weighted anatomical images were acquired with a rapid acquisition with relaxation enhancement (RARE) sequence (repetition time (TR)=5804.607s, effective echo time (TE)=32.5ms, echo spacing=10.83 ms, voxel size=0.125×0.125×0.5mm^3^, RARE factor=8, flip angle (FA)=90°). The BOLD-fMRI data were acquired by using a 2-D single-shot gradient echo echo-planar imaging (EPI) sequence (TR=1 s, TE=15 ms, FA=55°, in-plane resolution about 0.6×0.6 mm^2^, slice thickness=1 mm).

### Data preprocessing

MRI and fMRI data were preprocessed by using Analysis of Functional Neuroimages (AFNI) and custom-built functions in MATLAB. Within each session, the fMRI data were corrected for motion by registering every volume to the first volume using *3dvolreg*. After removing the first ten volumes, *retroicor* was used to correct for the motion artifacts due to respiratory and cardiac activity (32, 33). Then, *slicetimer* was used to correct the timing for each slice. For each animal, we first registered the EPI image to its T_2_ weighted structural images and then normalized to a volumetric template (34) using *flirt*. Motion artifacts were further corrected by regressing out the six motion-correction parameters. The fMRI data were then spatially smoothed with a 3-D Gaussian kernel with a 0.5-mm full width at half maximum (FWHM). The fMRI time series were detrended by regressing out a 2^rd^-order polynomial function voxel by voxel.

### General linear model analysis

We used the conventional general linear model (GLM) analysis to map the activations evoked by VNS as in previous studies (23, 24, 35-37). Specifically, we derived a response model by convolving the stimulation block (modeled as a box-car function) with a canonical hemodynamic response function (HRF) (modeled as a gamma function). For each session, the fMRI signal at every voxel was correlated with this response model. The correlation coefficient was converted to a z-score by using the Fisher’s z-transform. The voxel-wise z-score was averaged across sessions and animals, and the average z-score was evaluated for statistical significance with a one-sample t-test (p<0.05, uncorrected).

This analysis revealed the group-level activation map with VNS given an assumed response model. Since the validity of this response model for VNS was not established, we intentionally varied the response model by assuming three different values (3s, 6s, or 9s) for the peak latency of the HRF. We compared the activation maps obtained with the three different response models, to qualitatively assess the dependence of the model-based activation mapping on the presumed response characteristics.

### Independent component analysis

In contrast to the voxel-wise GLM analysis, we used ICA to map networks and their responses to VNS in a data-driven or model-free manner. For each session and each voxel, the fMRI signal during VNS was demeaned and divided by its standard deviation. The resulting fMRI data were then concatenated across all VNS-fMRI sessions. Infomax ICA (38) was used to decompose the concatenated data into 20 spatially independent components (ICs). Each of these ICs included a spatial map and a time series, representing a brain network and its temporal dynamics, respectively. In the spatial maps of individual ICs, the intensities at each given voxel represented the weights by which the time series of corresponding ICs were combined to explain this voxel’s fMRI time series. The polarity of each IC was determined to ensure the positive skewness of its weight distribution. Such weights were converted to Z-statistics and then thresholded as described in a previous paper (39). The threshold was selected such that the false negative rate was three times as large as the false positive rate. To obtain the false negative and positive rates, the Z-statistics of all voxels in an IC map were modeled as a two-Gaussian mixture distributions: one representing the noise, the other representing the signal.

Following ICA, we evaluated the VNS-evoked response separately for each IC, instead of each voxel. To do so, each IC’s time series was segmented according to the timing of every VNS block. Each segment lasted 54 seconds, starting from 3 seconds before the onset of a VNS block to 41 seconds after the offset of this block, while the stimulus block lasted 10 seconds. To address whether an IC responded to the VNS, we treated each time point as a random variable and each segment as an independent sample. One-way analysis of variance (ANOVA) was conducted against a null hypothesis that there was no difference among all the time points (meaning no response). The ICs that were statistically significant (*p*<5e-6) were considered as activated by VNS.

For each activated IC, we further characterized its temporal response to VNS. Briefly, we identified the time points during or after the VNS block, where the signals significantly differed from the pre-stimulus baseline by using the Tukey’s honest significant difference (HSD) test as a post-hoc analysis following the previous ANOVA test. Following this statistical test, the VNS-activated ICs were visually classified into three types (i.e. positive, negative, and mixed) of responses.

### Functional connectivity analysis

We further addressed whether VNS altered the patterns of temporal interactions among functional networks identified by ICA. To do so, the voxel time series was demeaned and standardized for each fMRI session including both VNS and resting conditions. The fMRI data were concatenated across all sessions and were then decomposed by ICA to yield 20 spatially ICs or networks, along with their corresponding time series. The first IC was removed because it was identified as the global component. The time series of the rest ICs were divided into the signals corresponding to the VNS sessions versus those corresponding to the resting-state sessions.

We defined the functional connectivity between networks as the temporal correlations between the ICs. The correlations were evaluated separately for the resting and VNS conditions and for every pair of ICs. Based on their temporal correlations, we grouped the ICs into clusters by applying k-means clustering method to ICs’ temporal correlation matrix. As a result, the correlations tended to be stronger within clusters than between clusters.

We further evaluated the differences in functional connectivity between the VNS condition and the resting state. For this purpose, the functional connectivity between ICs was evaluated for each VNS session, as well as each resting-state session. Their differences between these two conditions were evaluated using unpaired two-sample t-test (p<0.05, uncorrected). The changes in functional connectivity were displayed in the functional connectogram (40).

## Results

### Model-based VNS activations were sensitive to variation of the response model

Seven rats were scanned for fMRI while their left cervical vagus nerve was electrically stimulated in a (10s-ON-50s-OFF) block-design paradigm as illustrated in Fig 1. The BOLD response phase-locked to the VNS block appeared complex and variable across regions of interest (ROIs). For example, the BOLD responses were notably different across three ROIs, namely the retrosplenial cortex (RSC), brainstem (BS), and dorsal caudate putamen (Cpu) in a functional atlas (41) of the rat brain (Fig 2A). These responses were not readily explainable by a typical response model derived from a canonical HRF (Fig 2B, top). The GLM analysis with three different response models (by varying the HRF peak latency with 3s increments) yielded almost entirely distinctive activation maps (Fig 2B), each of which was only marginally significant (*p*<0.05, uncorrected). Therefore, VNS-evoked BOLD responses were too complex and variable to be captured by a single response model. The GLM analysis likely leads to incomplete and inconsistent activations with VNS, which possibly accounts for the diverging findings reported in the related literature (20-24).

**Fig 2.**
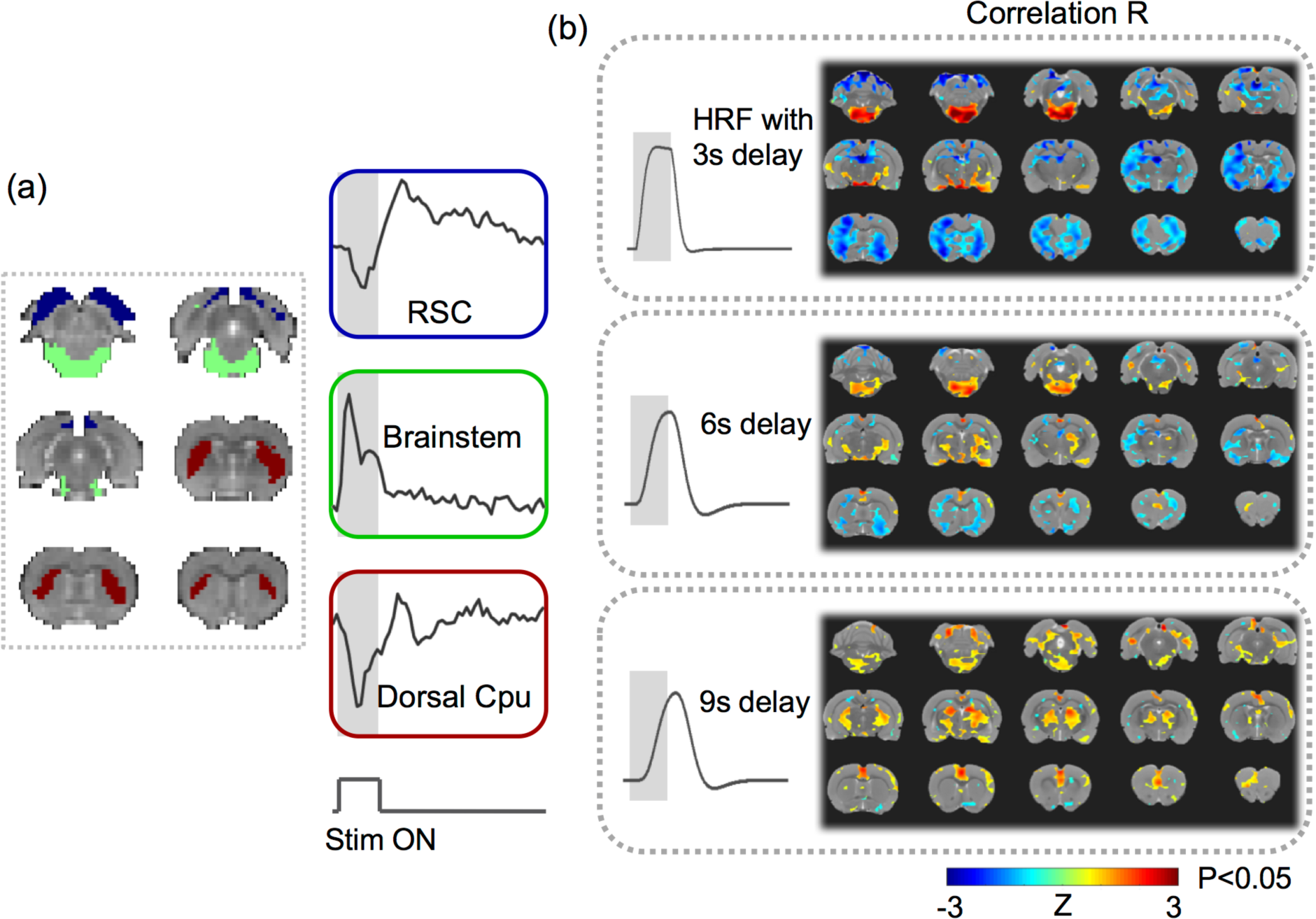
VNS-evoked responses varied across regions. (A) shows the response time series averaged within each of the three regions of interest: the retrosplenial cortex (RSC) (blue), the brainstem (green), and the dorsal caudate putamen (Cpu) (red). (B) shows the highly different activation maps based on the response models derived with the HRF, for which the peak latency was assumed to be 3s, 6s, or 9s. The color shows the group average of the z-transformed correlation between the voxel time series and the modeled response. The maps were thresholded with p<0.05 (one-sample t-test, uncorrected).

### VNS induced widespread and complex network responses

With a data-driven method, we evaluated the VNS-evoked responses in the level of networks, where the networks were identified as spatially ICs. It turned out that 15 out of the 20 ICs were significantly activated by VNS (one-way ANOVA, *p*<5e-6, Fig 3A). Those activated ICs collectively covered 76.03% of the brain volume (Fig 3B). Among the activated regions, the brainstem and the hypothalamus exhibited relatively stronger responses than other areas.

**Fig 3.**
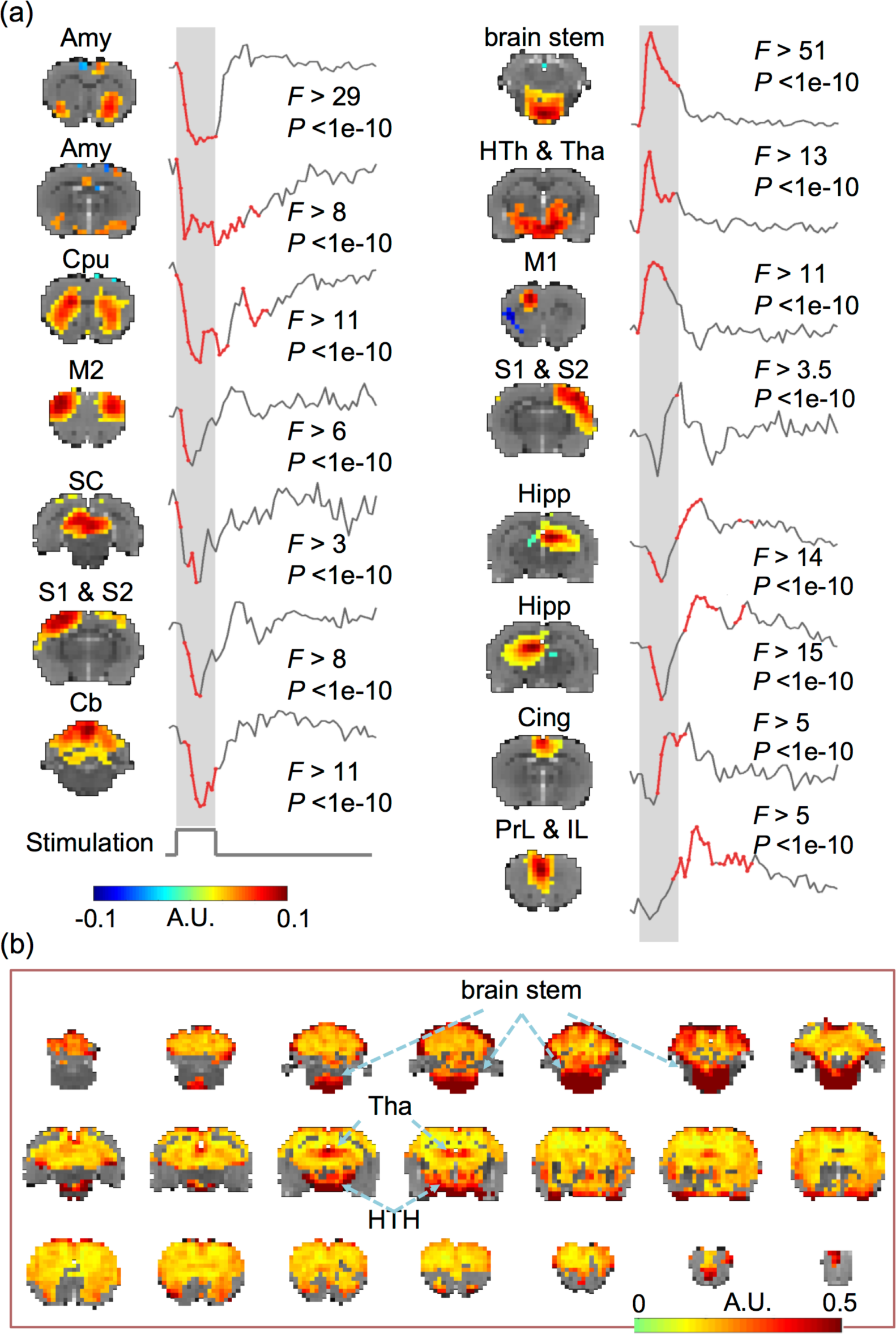
VNS evoked widespread and complex responses in the brain. (A) VNS-evoked responses for different brain networks derived with ICA. The ICA-defined networks are labeled as: amygdala (Amy), caudate putamen (Cpu), hippocampus, (Hipp), cingulate cortex (Cing), prelimbic cortex (PrL), infralimbic cortex (IL), brain stem, hypothalamus (HTh), thalamus (Tha), superior colliculus (SC), cerebellum (Cb), primary and secondary motor cortex (M1, M2), and primary and secondary somatosensory cortex (S1, S2). For each network, the time points at which the responses were significant are shown in red. (B) The VNS-activated voxels cover 76.03% of the brain volume. The color represents the standard deviation of the voxel-wise response averaged across repetitions of VNS. The locations with the greatest responses are highlighted with arrows. Data relevant to the VNS-evoked network responses are available in the online Supplementary Information.

The response time courses were also notably different across ICs. Fig 3A also highlights in red the time points, where the post-stimulus responses were significantly different from the pre-stimulus baseline (*p*<0.05, Tukey’s HSD). It was noticeable that different ICs were activated at different times following VNS. The response time courses also showed different polarities and shapes, and could be generally classified as the positive, negative, or biphasic-mixed response. The negative response was shown in the amygdala, dorsal striatum, primary motor cortex, midbrain, left somatosensory cortex, and superior cerebellum. The positive response was shown mainly in the brainstem, thalamus, and hypothalamus. The mixed response was shown in the hippocampal formation, cingulate cortex, and prelimbic & infralimbic cortex. The ICs that appeared to exhibit similar responses to VNS were presumably more functionally associated with one another. From a different perspective, the network-wise response to VNS also seemed to be either stimulus-locked or long-lasting (i.e. sustained even 20-30 s after the end of VNS). The stimulus-locked response was most notable in the brainstem and hypothalamus, which receives more direct vagal projections with fewer synapses. The long-lasting response was shown in the hippocampal formation, prefrontal cortex, amygdala, all of which were presumably related to high-level cognitive functions, such as memory formation, decision making, and emotion regulation. Speculatively, the former was the direct effect of VNS; the latter was the secondary effect.

### VNS altered functional connectivity

We further evaluated the network-network interactions during VNS in comparison with those in the resting state. The networks were captured as the ICs obtained by applying ICA to the data in both VNS and resting conditions. The matrix of pair-wise (IC-IC) correlations during VNS was overall similar to that in the resting state (Fig 4A). However, their differences in functional connectivity reorganized the clustering of individual networks (into Group 1, 2, 3) (Fig 4A). Group 1 covered the sensorimotor cortex, and it was mostly consistent between the VNS and resting conditions. Relative to the resting state, VNS reduced the extent of networks for Group 2, but increased the extent of networks for Group 3. For a closer investigation of the network reorganization, we found that VNS strengthened the correlations between the hippocampal formation and the retrosplenial cortex relative, but weakened the correlations between the prefrontal cortex and the basal ganglia. Beyond the difference in clustering, the significant changes in functional connectivity (*P*<0.005, *t*-test) are all shown in Fig 4B. The most notable changes were all related to the limbic system. During VNS, the cingulate cortex was less correlated with the ventral striatum; the hippocampal formation formed stronger functional connectivity between its left and right components, and with the retrosplenial cortex. The reorganization of functional connectivity was not only confined to the regions within the limbic system, but also between the limbic system and the sensorimotor cortex. VNS strengthened the interaction across the sensorimotor cortex with the hippocampal formation, retrosplenial cortex, and dorsal striatum, whereas it weakened the functional connectivity between the sensorimotor cortex and the cingulate cortex. In short, VNS reorganized the functional connectivity within the limbic system and altered the interactions between the limbic system and the sensorimotor cortex.

**Fig 4.**
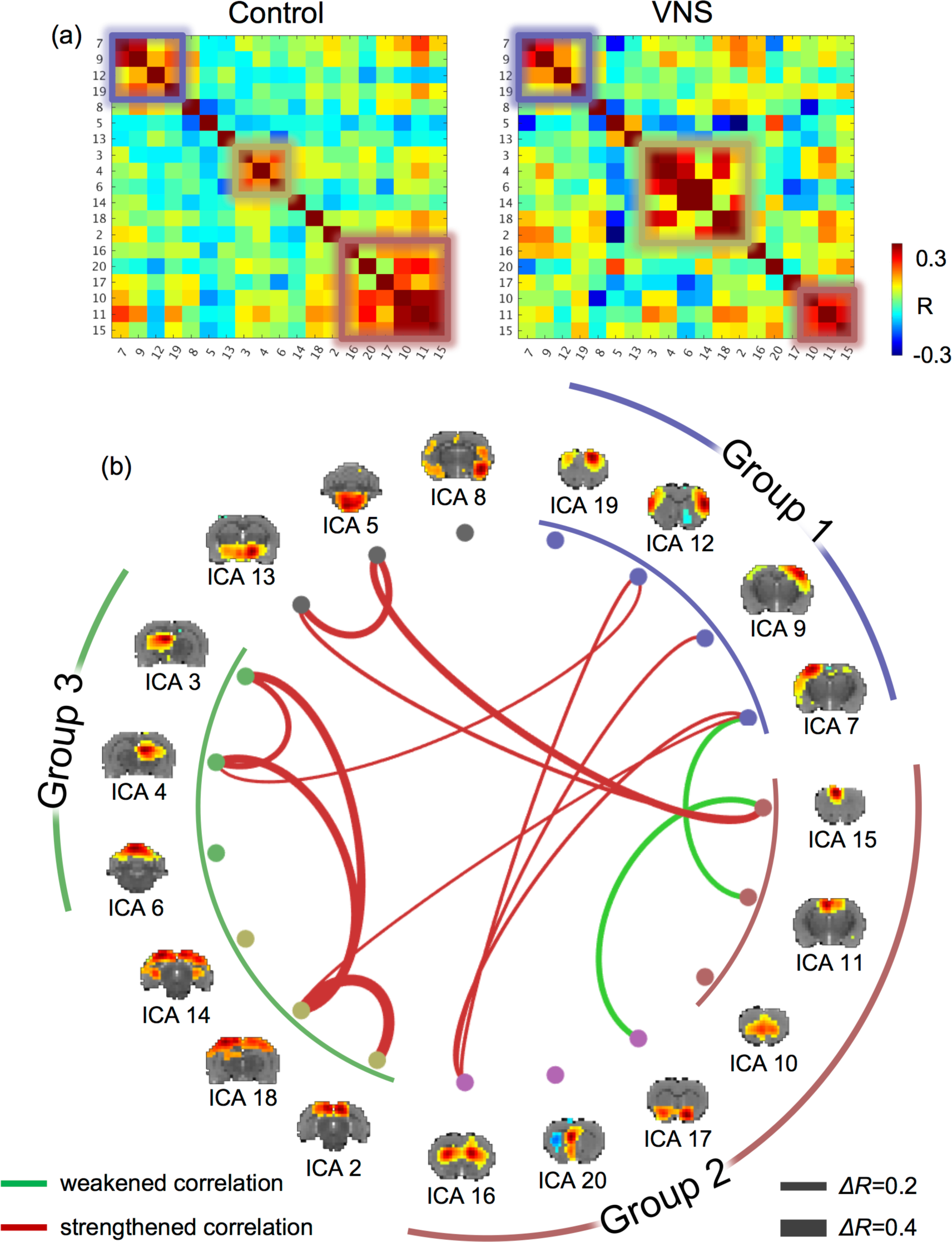
VNS altered the functional connectivity among functional networks. (A) shows the correlations between independent components. The left shows the correlation matrix during the resting state, or the “control” condition. The right shows the correlation matrix during VNS, or the “VNS” condition. Smaller squares highlight the networks (or ICs) that were clustered into groups (based on k-means clustering). (B) shows the IC-IC functional connectivity that was significantly different between the VNS and control conditions (t-test, P<0.005). Red lines represent increases in functional connectivity, and green lines represent decreases in functional connectivity. The thickness of the lines represents the (VNS minus control) change in correlation. The brain maps show the spatial patterns of individual ICs. Corresponding to the squares in (A), the arc lines illustrate how the ICs were clusters into groups, for the VNS condition (inner circle) and the control condition (outer circle).

## Discussion

Here, we report a model-free analysis method for mapping and characterizing the BOLD activations with VNS. Findings obtained with this method suggest that the repetitive and block-wise stimulation to the left cervical vagus nerve induces activations at widespread brain regions. The responses are complex and variable across regions, much beyond what can be described with conventionally assumed HRF. In addition, VNS also alters functional connectivity among different brain networks, and changes the brain’s functional organization from its intrinsic mode as observed in the resting state. These findings suggest widespread and profound effects of VNS on the brain’s regional activity and inter-regional interaction. Such effects are likely under-estimated by the model-based analysis in prior studies. This study also highlights the value of fMRI for addressing the large-scale and brain-wide effects of VNS, in order to understand and optimize its potential use for treatment of disease conditions in the brain or other organs, e.g. the gastrointestinal system.

### VNS evoke brain-wide responses

A major finding in this study was that VNS evoked time-locked and widespread BOLD responses over most parts of the brain. This finding appeared surprising at the first glance, since the simulation was applied to the left cervical vagus – a seemingly narrowly-focused entry of neuromodulation. Nevertheless, previous studies suggest that neural activity may drive global fluctuations in resting-state fMRI activity (42), and even simple (e.g., checkerboard) visual stimulation may evoke whole-brain fMRI responses (43). Common to those prior studies and this study is the notion that the brain is so densely wired and interconnected that focal modulation may induce a cascade of responses through neuronal circuits. Such network responses may even have a global reach, if the stimulation innervates sub-cortical structures with distributed modulatory effects on the brain (44).

In this regard, widespread responses to VNS may be mediated through the diffusive neuromodulation triggered by VNS. Vagal afferents project to the parabrachial nucleus, locus coeruleus, raphe nuclei through the nucleus of solitary tract (45). From the parabrachial nucleus, locus coeruleus, and raphe nuclei, connectivity extends onto the hypothalamus, thalamus, amygdala, anterior insular, infralimbic cortex, and lateral prefrontal cortex (46-49). In fact, the widespread VNS-evoked activations reported herein are consistent with the full picture gathered from piecemeal activations observed in prior VNS-fMRI (see reviews in (25)), transcutaneous VNS-fMRI (50), and EEG-fMRI studies (13, 14). In light of those results, the extent of the VNS effects on the brain has been under-estimated in prior studies, likely due to the use of simplified response models that fail to capture the complex and variable responses across all activated regions.

### Origins and interpretation of different response characteristics

Results in this study suggest that VNS induces a variety of BOLD responses that vary across regions. In addition to coarse and qualitative classification of various responses as positive, negative, or mixed (first negative and then positive) (Fig 3), the responses at various regions or networks also differed in terms of transient vs. sustained behaviors during and following VNS. For example, the responses at the brainstem and the hypothalamus showed a very rapid rise around 2s and rapid decay around 5s following the onset of VNS. Although the generalizable origins of transient BOLD responses are still debatable (51-53), we interpret the transient responses to VNS as a result of direct neuroelectric signaling through the vagal nerves. Nuclei in the brainstem, e.g., NTS, contain neurons receiving direct projections from the vagus, and in turn connect to the hypothalamus. Such brain structures are thus well-positioned to respond rapidly to VNS. Also, the rapid decay of the BOLD response in the brainstem and hypothalamus may indicate neuronal adaption – a factor of consideration for designing the duration and duty cycle of VNS. However, such interpretation should be taken with caution. The neurovascular coupling (modeled as the HRF) behaves as a low-pass filter through which the BOLD response is generated from local neuronal responses. Although the peak latency in HRF is 4 to 6s in humans, it is as short as 2s in rodents (54), making it relatively more suitable for tracking transient neuronal dynamics.

Another intriguing observation was the prolonged BOLD responses that sustained for a long period following the offset of VNS. In the striatum, hippocampus, as well as the prelimbic and infralimbic cortex, the VNS-evoked response lasted for 40s or even longer, while the VNS only lasted 10s (Fig 3). Such prolonged responses suggest potentially long-lasting effects of VNS. This observation is also in line with clinical studies showing that the effects of VNS on seizure suppression are not limited to the stimulation, but sustain during periods in the absence of VNS (55). Moreover, those regions showing prolonged effects of VNS tended to be higher-level functional areas presumably involved in learning, decision-making, memory, and emotion-processing. Speculatively, it implies that the VNS-based modulation of cognitive functions or dysfunctions operates in a relatively longer time scale while imposing potentially therapeutic effects on a longer term.

### VNS alters network-network interactions

This study highlights the importance of evaluating the effects of VNS on functional connectivity, which measures the degree to which regions or networks interact with each other. It is widely recognized that brain functions emerge from coordination among regions (56). However, prior studies address the effects of VNS in focal and target regions (57), whereas the effects of VNS on functional connectivity is perhaps more functionally relevant. In line with this perspective, a recent study has shown that transcutaneous VNS modulates the functional connectivity in the default mode network in patients with major depressive disorders, and the change in functional connectivity is related to the therapeutic efficacy across individual patients [50]. Thus, VNS may reorganize the patterns of interactions among functional networks – a plausible network basis underlying VNS-based therapeutics.

Our results show that VNS reorganizes the functional connectivity with respect to the limbic system. Relative to the intrinsic functional connectivity in the resting state, VNS increases functional connectivity between the retrosplenial cortex and hippocampal formation, of which the functional roles are presumably related to memory, learning, and monitoring sensory inputs (58, 59). In addition, VNS increases functional connectivity between the sensory cortex and the striatum, of which the functional roles are presumably the integration of sensorimotor, cognitive, and motivation/emotion (60). In contrast, VNS decreases functional connectivity between the cingulate cortex and the ventral striatum, which likely affect the emotional control of visceral, skeletal, and endocrine outflow (61). Collectively, these observations lead us to speculate that VNS biases the limbic system to shift its functional role from emotional processing to perceptual learning. Such speculation is consistent with the therapeutic effects of VNS in depression patients (62, 63) and the cortical plasticity of interest to perceptual learning and motor rehabilitation induced by VNS (64, 65).

### Model-free activation mapping in the level of networks

Central to this study is the use of model-free and data-driven analysis for mapping activations in the level of networks, instead of voxels or regions. This is in contrast to conventional GLM analysis used in previous VNS-fMRI studies, which assumes that neural responses sustain in the entire period of VNS, and the BOLD effects of neural responses may be modeled with a canonical HRF (23, 24, 35-37). Both of these assumptions may not be entirely valid. Neural responses may exhibit a range of non-linear characteristics. The typical HRF model is mostly based on data or findings obtained from the cortex during sensory stimulation (66), whereas no study has modeled the HRF for VNS. Moreover, the neurovascular coupling may also vary across regions in the brain, especially between sub-cortical and cortical areas due to their differences in local vasculatures (67). Thus, the GLM analysis with a single and empirical response model most likely falls short for explaining the complex and variable responses across all brain regions.

The model-free analysis allowed us to detect the VNS-evoked brain response in a data-driven way without assuming any prior response model. A similar strategy has been used to test for voxel-wise BOLD responses to visual stimulation in humans (43). What was perhaps unique in this study was the use of the model-free analysis on the activity of spatially independent components, rather than that of single voxels. Each IC contained a set of voxels (or locations) that shared a common pattern of temporal dynamics. ICA utilized the fact that individual voxels were organized by networks, not in isolation, to extract the network activity as the time series of each IC, which reflected the (weighted) average activity of all the voxels that belonged to each IC. As such, the signal-to-noise ratio was higher for IC-wise activity than for voxel-wise activity, providing better sensitivity for detecting activations at the network level. This model-free analysis method is thus arguably more favorable than conventional GLM analysis, especially when the response characteristics are complex and unclear, e.g., given VNS.

### Potentially confounding cardiac and respiratory effects

The BOLD signal is an indirect measure of neural activity. Therefore, it may be affected by non-neuronal physiological fluctuations (68). Previous studies have shown that VNS causes cardiorespiratory effects, e.g., variations in the heart rate, respiration rate, and SpO2 (69, 70). Such effects may potentially confound the interpretation of the VNS-induced BOLD responses regarding neuronal activations. Such confounding effects were highlighted in a prior study, in which VNS was found to decrease the heart rate and in turn decrease the BOLD signal throughout the rat brain (71). In this study, we were concerned about this potential confounding effect, and reduced to the pulse width of VNS to 0.1ms, instead of 0.5ms in that study (71). Such shortened pulse width largely mitigated the cardiac effects, as no obvious changes in heart rate were noticeable during experiments.

Also, the cardiac effects would manifest themselves as the common physiological response observable throughout the brain. This was not the case in this study. Despite wide-spread activations with VNS, the responses at individual regions exhibited different temporal characteristics, which could not be readily explained by a common confounding source (e.g., respiratory or cardiac). Instead, the regions with similar response patterns formed well-patterned functional networks, resembling intrinsic resting-state networks previously observed in rats (72). For these reasons, it was unlikely that the VNS-induced activation and functional-connectivity patterns were the result at non-neuronal cardiac or respiratory effects. Similar justifications are also applicable to the confounding respiratory effects. Nevertheless, future studies are desirable to fully disentangle the neuronal vs. non-neuronal effects of VNS. Of particular interest is using multi-echo fMRI to differentiate BOLD vs. non-BOLD effects (68), and combining electrophysiology and fMRI to pinpoint the neural origin of the fMRI response to VNS (73).

## Acknowledgements

The authors thank Elizabeth Baronowsky and Dr. Matthew Ward for their initial assistance in animal surgery. This work is supported by the National Institutes of Health [OT2TR001965, R01MH104402].

## Supplementary Information

ICA-derived network responses to VNS, including the spatial patterns of the networks, and their corresponding response time courses.

